# TLCellClassifier: Machine Learning Based Cell Classification for Bright-Field Time-Lapse Images

**DOI:** 10.1101/2024.06.11.598552

**Authors:** Qibing Jiang, Praneeth Reddy Sudalagunta, Mark Meads, Xiaohong Zhao, Alexandra Achille, David Noyes, Maria Silva, Rafael Renatino Canevarolo, Ken Shain, Ariosto Silva, Wei Zhang

## Abstract

Immunotherapies have shown promising results in treating patients with hematological malignancies like multiple myeloma, which is an incurable but treatable bone marrow-resident plasma cell cancer. Choosing the most efficacious treatment for a patient remains a challenge in such cancers. However, pre-clinical assays involving patient-derived tumor cells co-cultured in an *ex vivo* reconstruction of immune-tumor micro-environment have gained considerable notoriety over the past decade. Such assays can characterize a patient’s response to several therapeutic agents including immunotherapies in a high-throughput manner, where bright-field images of tumor (target) cells interacting with effector cells (T cells, Natural Killer (NK) cells, and macrophages) are captured once every 30 minutes for upto six days. Cell detection, tracking, and classification of thousands of cells of two or more types in each frame is bound to test the limits of some of the most advanced computer vision tools developed to date and requires a specialized approach. We propose TLCellClassifier (time-lapse cell classifier) for live cell detection, cell tracking, and cell type classification, with enhanced accuracy and efficiency obtained by integrating convolutional neural networks (CNN), metric learning, and long short-term memory (LSTM) networks, respectively. State-of-the-art computer vision software like KTH-SE and YOLOv8 are compared with TLCellClassifier, which shows improved accuracy in detection (CNN) and tracking (metric learning). A two-stage LSTM-based cell type classification method is implemented to distinguish between multiple myeloma (tumor/target) cells and macrophages/monocytes (immune/effector cells). Validation of cell type classification was done both using synthetic datasets and *ex vivo* experiments involving patient-derived tumor/immune cells.

**Availability and implementation:** https://github.com/QibingJiang/cell classification ml

## Introduction

Immunotherapy [1] is a type of cancer treatment that utilizes the body’s immune system to combat cancer [2, 3]. It can enhance or modify the immune system’s functionality, enabling it to locate and attack cancer cells [1, 4]. Immunotherapy may be administered alone or in conjunction with other cancer treatments [5, 6]. Preclinical *ex vivo* assays [7, 8, 9, 10] can help identify the most efficacious therapy for each patient by treating co-cultures of tumor and immune cells with several drugs/combinations in a reconstruction of the tumor microenvironment. The use of bright-field imaging for high-throughput drug sensitivity characterization facilitates quantifying *ex vivo* responses in a temporal fashion unlike traditional end-point assays. These bright-field images typically feature thousands of cells in each frame and are captured once every 30 minutes for up to six days. While such high-throughput time-lapse imaging of *ex vivo* cultures yield high fidelity data and provide a unique insight into the dynamic nature of patient/drug-specific *ex vivo* responses, they pose a significant challenge in accurately detecting and tracking [11, 12] thousands of cells across hundreds of frames. Additionally, co-culture assays involve multiple types of cells and require an efficient approach to quantify the targeted effect of the therapy on the tumor cells compared to other non-malignant cellular subtypes [13], which presents a unique challenge to classify cell types based on morphology and behavior. In this article, we systematically address three main challenges, namely, detection of cancer cells and identifying cell-specific features, tracking individual cells across time using time-lapse imaging, and classifying cells into their respective cell types.

Images of cells can be captured using various imaging modalities, including 2D bright-field, phase contrast [14], differential interference contrast (DIC) [15], and fluorescence images [16]. Fluorescence images are widely used by biologists due to the rich details they provide in cells. Sequences of stained or counterstained cells or nuclei moving on a flat substrate can be captured as 2D or 3D time-lapse sequences, offering additional insights into the environment and behavior of the cells. The Cell Tracking Challenge [17] contains various datasets and many state-of-the-art methods [18, 19, 12] have been developed to address detection and tracking challenges in various scenarios. For both cell detection and tracking, numerous deep learning-based methods and canonical approaches have been proposed, where thresholding and learning-based methods are effective in segmenting DIC images [20]. Moreover, thresholding-based pipelines and level-set-based approaches were proposed to detect cells in bright-field microscopy images [21, 22]. Over the past decade, deep learning methods have progressively outperformed canonical methods in detection tasks. For instance, DKFZ-GE, one of the methods in the Cell Tracking Challenge, employs nnU-Net [23] and achieves the best performance on the Fluo-C3DH-A549 dataset. U-net [24] based methods perform well on general object detection and medical image segmentation, which is made possible by high resolution images. Another method from Cell Tracking Challenge, UNSW-AU utilizes NAS (neural architecture search) [25] and performs very well on the Fluo-C2DL-MSC dataset. Advanced machine learning methods automatically extract numerous features in an unsupervised manner, surpassing the efficacy of handcrafted features such as setting thresholds for pixels. However, compared to cell detection, deep learning methods have not shown significant improvements in cell tracking.

Most tracking methods fall into one of two categories. The first relies on tracking overlapping objects from two consecutive images [26, 27, 28, 29]. This approach is particularly effective for large, slow-moving cells captured using high-sampling-rate imaging systems, ensuring an overlap for each cell in two consecutive images. The second class of methods is based on Euclidean distance. A temporal association is established between sets of cells detected in different images. This association can be built either on two consecutive images or on a multi-image sliding window that includes all images simultaneously. The Hungarian algorithm [30] has been utilized to align cells on two sequential images, as demonstrated in studies [31, 32]. On the other hand, the Viterbi algorithm can be employed to match cells across all images and establish global cell tracks, as shown in the KTH-SE method [33]. Additionally, some models utilize probabilistic approaches to build temporal links, as indicated in studies [34, 35]. However, predicting cell trajectories remains a significant challenge, particularly in cases where cell migration resembles Brownian motion.

Cell classification is also a critical area of research in medical image processing. For instance, Inception v3 and artificial features have been combined to classify cervical cell images in the HErlev Pap Smear Dataset [36]. Similarly, CNN-based methods are employed to classify white blood cells in blood smear images [37], and a combination of CNN and support vector machine is used to classify single-cell videos into patterns and impurities [38]. Additionally, deep neural networks have been utilized to rapidly detect live bacterial growth and classify corresponding species in time-lapse images [39]. These studies typically involve datasets with high-resolution images of single cells or videos capturing dozens of images per second. However, cells on smear slides have limited viability, restricting cell motion and evolution. Our dataset is unique because it monitors thousands of cells in a liquid medium for six days. This extended observation period requires the development of a novel computer vision pipeline capable of classifying thousands of cells in these new *ex vivo* conditions.

Most computer vision software discussed relies on high-resolution images or images stained with fluorescent markers, which works well on images of fixed cells such as FFPE (Formalin fixed Paraffin Embedded) tissue slides. However, fluorescence imaging of live cells suffer from drawbacks such as photobleaching and phototoxicity [40]. This makes bright-field imaging ideal for capturing cell membrane motion, cell morphology, and motility in time lapse imaging. We are working with a dataset [8, 9, 10] developed at Moffitt Cancer Center that involves bright-field images of patient-derived CD138+ multiple myeloma (MM) cells co-cultured with CD14+ macrophages/monocytes along with human bone marrow stromal cells captured once every 30 minutes for up to six days. Each 1328 *×* 1048 bright-field image contains thousands of cells, and the per cell resolution is very low and often appears pixelated. Unlike the images typically studied in contemporary research, which typically contain only dozens of cells, our dataset’s larger cell population makes cell tracking challenging due to an exponentially growing execution time with increased cell number [41]. CNN-based methods [42] have demonstrated significant capabilities in image processing tasks such as object detection, tracking, and classification. Notable examples include ResNet [43], YOLO [44], and GoogleNet [45]. However, these methods excel when per cell resolution is high [46]. A key challenge with CNN-based methods is that as the feature map progressively reduces in size when traversing deeper into the CNN layer, smaller objects tend to vanish in the deeper layers, making their detection and tracking particularly challenging [47, 48, 49].

This article explores the detection and tracking of patient-derived MM cells in bright-field time-lapse images derived from an *ex vivo* assay. These images are captured to screen effective drugs for MM cells. In each experiment, MM cells derived from one patient are co-cultured with human bone marrow stromal cells with either macrophages/monocytes, NK cells, and T cells in 384 wells, each containing different drugs/combinations at five serially diluted concentrations. By capturing images of each well for six days, we can observe the dynamic interactions of the tumor with its microenvironment, immune effector cells, and therapeutic agents. To quantify the viability of the MM cells, we have developed an image processing pipeline TLCellClassifier (time-lapse cell classifier) that includes CNN-based detection, metric learning-based tracking [50], and long short-term memory (LSTM)-based [51] classification methods. These methods are described in detail in Methods. The evaluation of the proposed method and its comparison with baseline approaches are explored in the Results section. Finally, we conclude our work and discuss future directions.

## Methods

Time-lapse images utilized in this study comprise bright-field images of MM cells captured under a microscope. These bright-field time-lapse images are obtained through the *ex vivo* assay mentioned below and designed by our collaborators [7, 8, 9, 10]. Fig.1 illustrates the overall workflow of the proposed framework, which consists of three main components: cell detection, tracking, and classification. Cells captured in bright-field images can be distinguished from the background via a sharp gradient in contrast. However, the presence of various cellular subtypes of different sizes and shapes that are captured at different brightness intensities can complicate the problem of cell detection. To address this, a CNN-based cell detection approach is developed. Moreover, the crowded nature of cells and their similar appearances present tracking difficulties. In this framework, metric learning is utilized to track cells based on their neighborhoods. Lastly, cells are classified into three types: macrophages, MM, and others (with CD14+ expression but not macrophages, such as monocytes). LSTM is applied here to classify single-cell tracks/videos. The details of the three components are described in the following subsections, followed by the introduction of the baseline and benchmark methods.

**Fig. 1.**
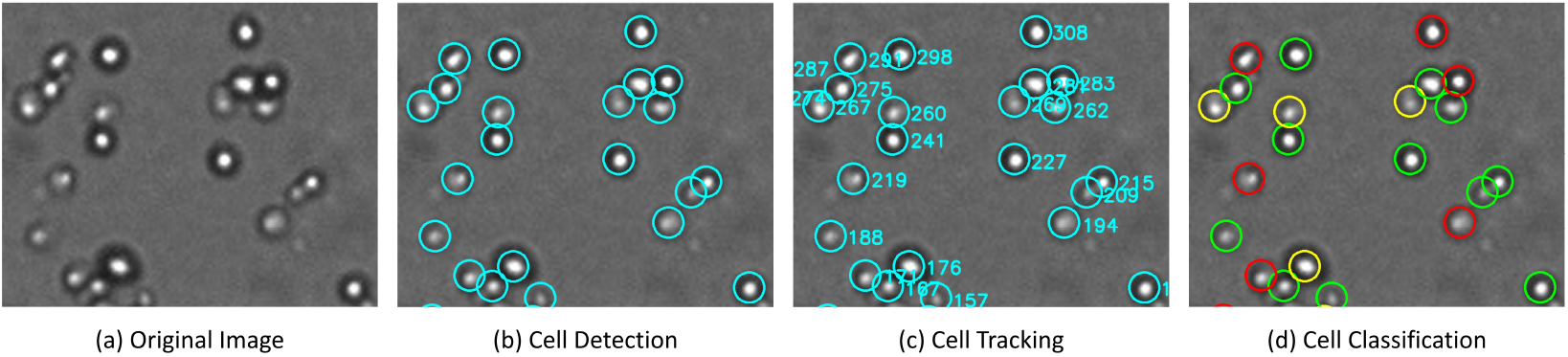
The overall workflow of the pipeline is as follows: (a) Some cells appear bright while others appear dark in the bright-field images. (b) A CNN-based cell detection method is designed in this research. (c) Due to the crowded and similar appearance of cells, metric learning is applied in this framework to track cells based on their neighborhoods. (d) Cells are classified into three types: Macrophage (highlighted in red), MM (highlighted in green), and others (highlighted in yellow). LSTM is then applied to classify the cells.

### *Ex V ivo* Assay

To evaluate the impact of drugs on primary MM cells, our collaborators at Moffitt Cancer Center conduct an *ex vivo* assay [8]. In this assay, primary MM cells, along with macrophage and stromal cells, are isolated from fresh bone marrow samples of patients. These cells are then separated and plated into 384-well plates, providing a supportive microenvironment for their survival. Various drugs, each at five different concentrations, are added to designated wells, while ten control wells remain untreated. The efficacy of a drug for MM treatment is determined by observing the rate of MM cell death in drug-treated wells compared to control wells. Additionally, differences in MM cell viability across different drug concentrations indicate concentration-dependent effects of the drug on MM cells. Conversely, if there is no variation in MM cell viability among wells with different drug concentrations, it suggests that varying concentrations have similar effects on MM cells. The plated cells are observed using a motorized stage microscope, maintaining conditions similar to those in the human body. Images at a resolution of 1328 *×* 1048 pixels are captured under bright-field microscopy every half an hour for up to six days. In these images, macrophage and MM cells appear as white nearly circular objects, while stromal cells are observed as white elongated structures, as shown in Fig.2. Fig.2 also shows a more challenging case: ‘Myeloma 2’, which is darker than the background. These kinds of images are captured when a particular well is slightly out of focus.

**Fig. 2.**
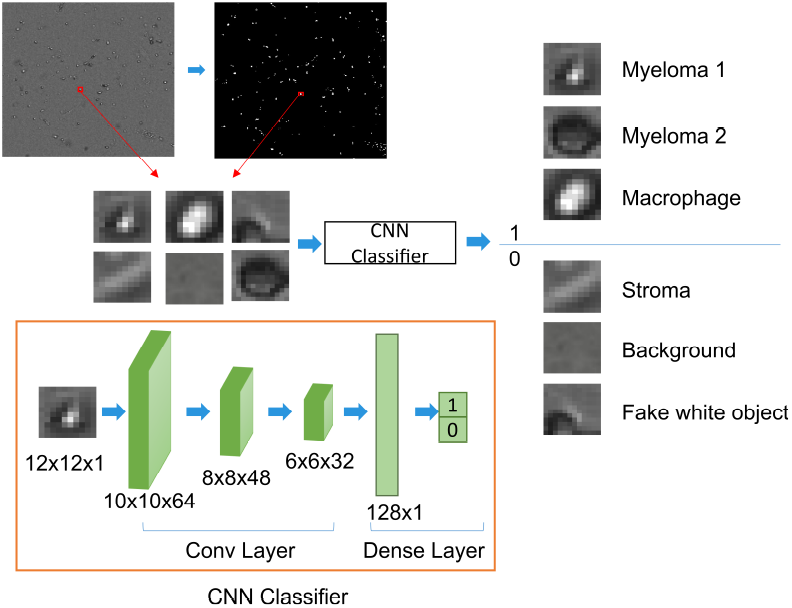
The image is initially converted into a binary image. For each cluster of white pixels detected in the binary image, an ROI is cropped from the original image. These ROIs are then classified by a CNN classifier into two categories: 1. ROUND CELL and 2. NEGATIVE OBJECT.

### Cell Detection

Both MM cells and macrophage cells appear as white objects in the bright-field images, varying in size and shape. A bounding box around each cell can typically have a resolution as low as 12*×*12 pixels. In a CNN-based method, smaller objects may vanish as the image passes through deeper layers. To address this, we integrated traditional image processing techniques (thresholding) with a shallow CNN for cell detection. Cells are detected in two steps, as illustrated in Fig.2. Initially, the grayscale image is converted into a binary image using a predefined threshold [12]. The bounding box for each candidate cell, a 12*×*12 region of interest (ROI), is then cropped from the grayscale image. These ROIs serve as input for the CNN classifier, which categorizes them into two classes: ROUND CELL, encompassing macrophage and MM cells, and NEGATIVE OBJECT, which includes stroma, background, and other white flares.

The grouping of both macrophages and MM cells into the same category is a result of the limitations of human visual perception in distinguishing these two cell types within a single image. However, macrophage and MM cells exhibit distinct behaviors over time, making it possible to differentiate them through time-lapse video analysis. The following section explores video classification techniques aimed at separating these two cell types. Within this section, macrophages, MM, and other immune cells are initially detected and categorized as ROUND CELL.

From Fig.2, we can observe that the first category, ROUND CELL, contains cell images that appear different. Cell images can vary due to many factors. Maintaining consistent focus on cells while capturing thousands of wells across different plates is challenging. Additionally, cells can reside at different depths within the culture medium, and the well-plate may have a slight curvature, all of which contribute to the variations observed in MM cells, as shown in Fig.2. To enhance the robustness of the CNN classifier, a diverse range of MM images is collected in the training dataset.

Our CNN classifier consists of four layers, including three ResNet convolution layers and one dense layer. The CNN architecture is kept shallow due to the small size of cells. Rectified Linear Unit (ReLU) is utilized as the activation function in the convolutional layers, and Softmax is employed in the dense layer. The loss function is defined as follows:

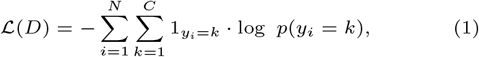

where *N* is the number of cell images and *C* is the number of classes, which is two (1. ROUND CELL, 2. NEGATIVE OBJECT) in this study. *y*_*i*_ represents the true label (ground truth) for the (i)-th cell image, while *p*(*y*_*i*_ = *k*) denotes the predicted probability that the (i)-th cell image belongs to class *k*. 1_*y*=*k*_ is an indicator function, returning 1 when *y* equals *k*, and 0 otherwise. The predicted probability is generated by the final layer of the CNN. The process of minimizing the cross-entropy loss selects parameters that drive the predicted probability towards 1 for the actual class.

For each image, cell detection is accomplished in two steps: ROI identification and classification. This process yields cell coordinates, contours, eccentricity, and other relevant information. Additionally, a 12×12 image is cropped for each cell, enabling further analyses such as tracking and cell type classification.

### Cell Tracking

Using the aforementioned cell detection approach, ROIs in each frame across time are identified. To create cell tracks for each cell across time, the detected cells within the ROIs must be matched. Tracking cells presents unique challenges compared to tracking objects in daily life, as cells often appear similar and are densely packed. Drawing insights from person identification techniques [52, 53], we discovered that metric learning can be trained to generate embedding vectors capable of representing the similarity or dissimilarity features between two images. The challenge in our dataset lies in the similarity of MM cells, which are difficult to distinguish due to low resolution. While different persons can be distinguished based on appearance, it is not always possible to differentiate MM cells with a 12×12 resolution. Through observation of time-lapse images, we found that the cell neighborhood remains relatively stable and recognizable between two time points. Therefore, instead of only tracking individual cells, we also track cell neighborhoods, as demonstrated in Fig.3.

**Fig. 3.**
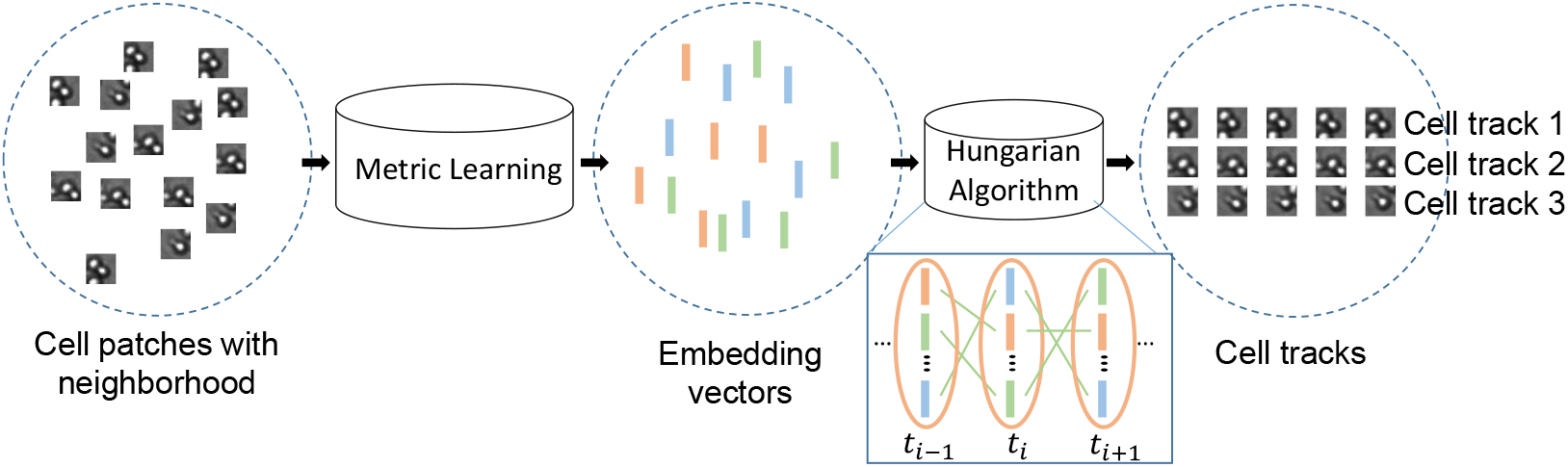
Framework for cell tracking. Cell neighborhood images are the input of the metric learning-based model, which generates embedding vectors as output. The Hungarian algorithm is then utilized to match the vectors obtained from two different time points. By matching these embedding vectors, cells from the two time points can be effectively matched.

Our metric learning model is designed for object re-identification in images captured at different time points. The architecture of the metric learning model, as summarized in Table 1, uses larger ROIs of size 18*×*18 pixels. These images consist of a 12*×*12 ROI detected previously along with its surrounding area. Through a series of convolutional layers, the image dimensions are reduced to 2*×*2 pixels. Subsequently, an embedding vector of length 128 is extracted using a dense layer.

**Table 1.**
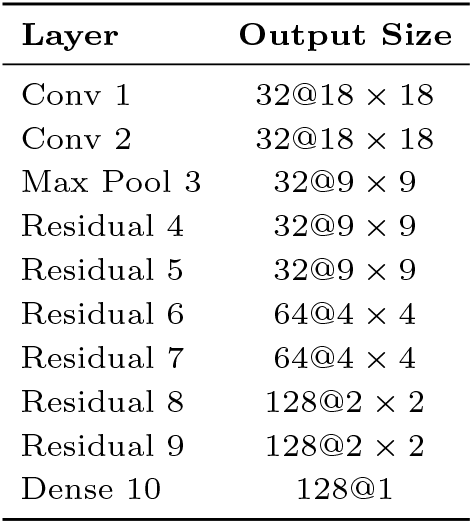
Architecture of the metric learning-based model

| Layer | Output Size |
| --- | --- |
| Conv 1 | $32@18 \times 18$ |
| Conv 2 | $32@18 \times 18$ |
| Max Pool 3 | $32@9 \times 9$ |
| Residual 4 | $32@9 \times 9$ |
| Residual 5 | $32@9 \times 9$ |
| Residual 6 | $64@4 \times 4$ |
| Residual 7 | $64@4 \times 4$ |
| Residual 8 | $128@2 \times 2$ |
| Residual 9 | $128@2 \times 2$ |
| Dense 10 | $128@1$ |

To enhance accuracy, the network incorporates several residual blocks [43], enabling information flow from the initial layers to the final ones.

The loss function utilized for training enforces separation based on the embedding vectors of images within individual cell tracks, ensuring they differ from the mean of the embedding vectors across other cell tracks:

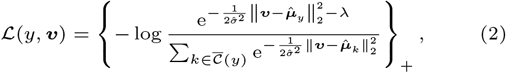

where ***v*** represents one embedding vector (one input image) to be learned in cell track 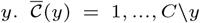 denotes the set of cell tracks excluding *y. λ* is a hyper-parameter, 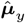 is the mean of the cell image predictions in cell track *y*, and 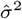 is the variance of all cell images. By minimizing the loss, the feature representation from the same cell will become closer, while the feature representation from different cell tracks will become further apart.

Another challenge in tracking cells arises from the presence of macrophages in our time-lapse images, which move rapidly and irregularly. Unlike other cells, macrophages do not maintain a stable neighborhood, making it difficult for metric learning to match all detected cells between two time points. Therefore, an additional step is needed to track these cells, involving the calculation and comparison of cell locations between two time points to match them.

The overall tracking algorithm is outlined in Algorithm 1, where 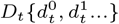 and 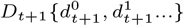 represent cell detections at time points *t* and *t* + 1, respectively. MLM and HA denote the Metric Learning Model and Hungarian Algorithm, respectively. Cell tracks are established by matching detected cells from two consecutive time points. For each pair of adjacent time points, the matching process consists of two steps. In the first step (lines 8-19), the metric learning-based model generates embedding vectors *V*_*t*_ and *V*_*t*+1_ based on the neighborhood of each cell. Then, the cosine similarity between the vectors from the two time points is calculated. Finally, the Hungarian algorithm is used to match the embedding vectors. Only pairs with a cosine similarity score *score*(*a*) higher than the threshold are retained (line 11). For each matched pair 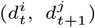, the current detection 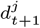 is added to the track *T* [*k*] that includes 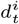 (line 13). Fast-moving cells, which lack a stable neighborhood, may not find a match in the first step (line 14). This is where the second step comes into effect (lines 20-23). First, the Euclidean distance is calculated between the coordinates of the cells from the two time points. Then, the Hungarian algorithm is applied to match the cells. After completing both steps, cells from the two time points are matched. By applying these steps for each pair of consecutive time points, cell tracks are established.

### Cell Classification

Cell classification plays an important role in drug screening. Within the microenvironment of a well, various cell types such as macrophages, MM cells, and others coexist. As the MM cells are treated with drugs, some cells undergo apoptosis, while others may be subject to phagocytosis over time. It is important to know the proportion of each cell type and monitor the number of cells of each type that undergo cell death. In this study, our primary objective is to identify a drug capable of selectively targeting MM cells while avoiding any detrimental effects on immune cells.

#### Algorithm 1

Tracking Algorithm

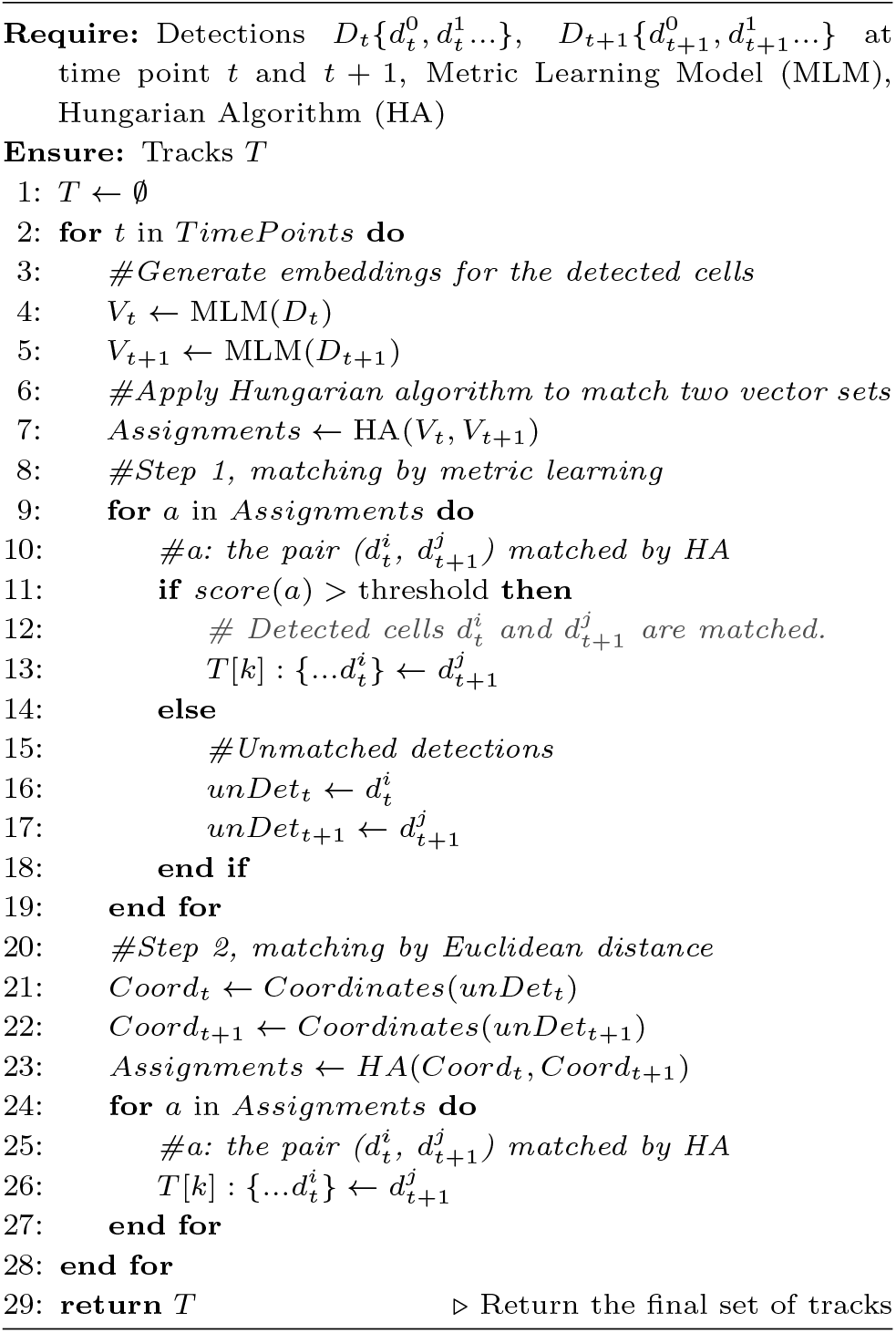

Based on cell detection and tracking, we can crop single-cell images at each time point and track for each cell. With thousands of single-cell tracks available, cells can be classified into different types: myeloma, macrophage, or others. Although these cells may look similar in a single image, their behavior differs across time points. Therefore, LSTM is employed to classify the cells by analyzing the single-cell tracks. The architecture of our LSTM method is illustrated in Fig.4.

The CNN model in Fig.4(a) is a pre-trained Inception v3 [54]. Images of 1000 different classes in the ImageNet [55] are used to train Inception v3. The Inception v3 architecture is a deep CNN used for image analysis and object detection. It is designed to allow deeper networks while also keeping the number of parameters from growing too large. The architecture is made up of multiple Inception modules, which consist of parallel layers of convolutions and pooling. The Inception v3 architecture has been employed in many different applications. In this study, it is utilized to generate feature representations that serve as the input for the LSTM module.

The input to the Inception v3 model is a 12×12 cell image in a single-cell track. The output of the model is a feature vector containing extracted information from the cell image. Inception v3 is followed by the LSTM model for multi-class classification. Categorical cross-entropy is used as the loss function, which is the same as Equation (1). In this classification task, *N* represents the number of cell tracks in the dataset, and *C* represents the number of classes, including macrophage, myeloma, and ‘others.’ The objective is to optimize the LSTM model parameters to minimize this loss.

Monocytes, categorized under ‘others,’ are white blood cells that differentiate into macrophages. In this precursor state, these cells have a different morphology and behavior. In *ex vivo* cultures, the differences between monocytes and myeloma are difficult to distinguish. Even through the observation of single-cell tracks, it remains difficult to separate these two cell types. To compel the model to learn specific features for each type, two stages of LSTM are employed. In the first stage, the LSTM-based Classifier 1 is tasked with separating macrophages from other cells, while in the second stage, the LSTM-based Classifier 2 distinguishes between other cells (such as monocytes) and myeloma, as illustrated in Fig. 4(b).

**Fig. 4.**
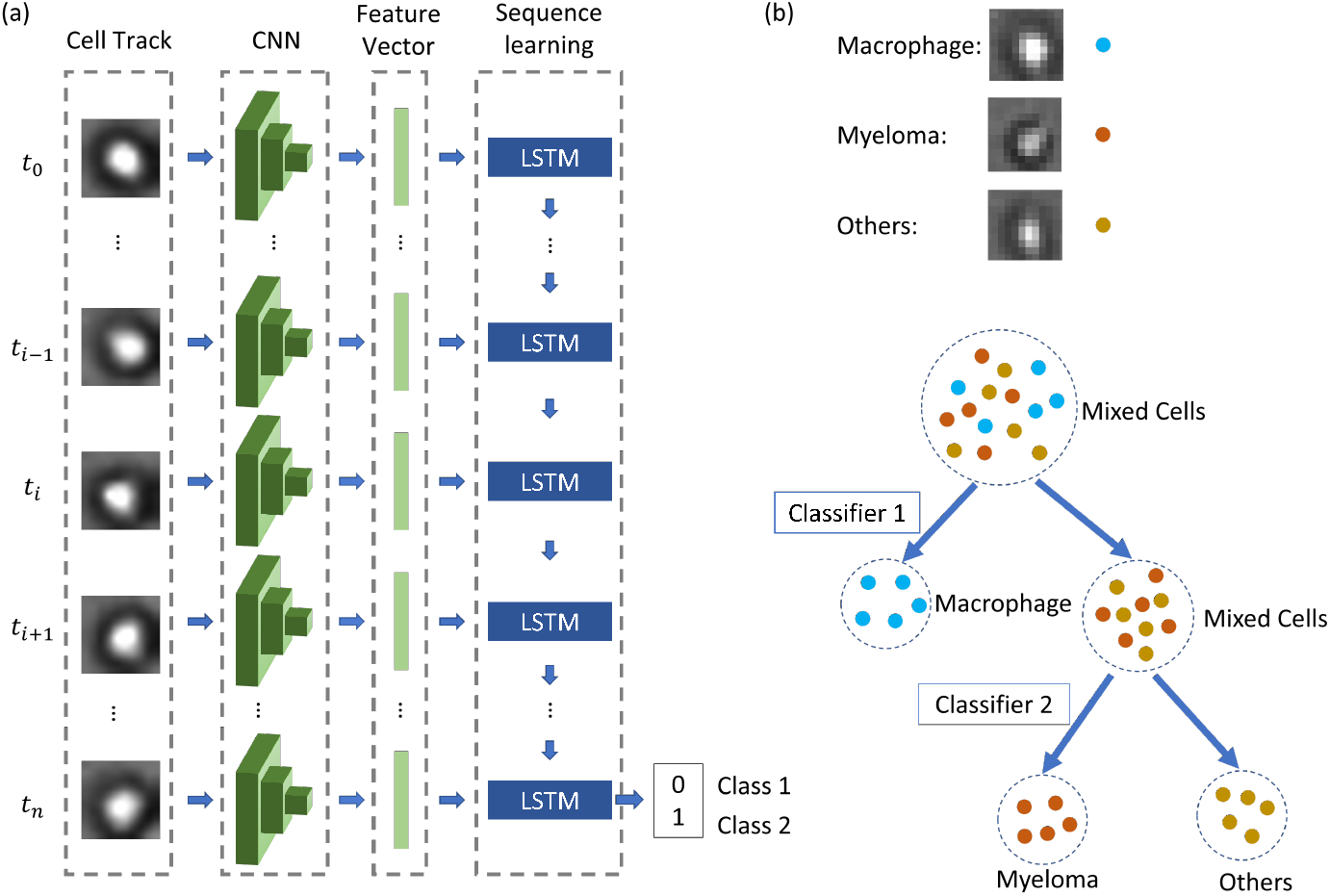
LSTM-based Cell Classification. (a) Inception V3 is employed to generate a feature vector for each cell image, which is then utilized by LSTM for cell classification. (b) Two stages of LSTM are designed in this approach. The first stage aims to classify cells into either *{*myeloma, others*}* or *{*macrophage*}*. The second stage further refines the classification, distinguishing between myeloma and other cell types.

To train the two LSTM-based classifiers, we prepared two types of experiments (i.e., wells). The first contains only MM cells. The second consists of immune cell-only wells, which contain only CD14+ selected cells (macrophages/monocytes). **(1)** Classifier 1 is trained with the dataset of the single-cell tracks generated from the two types of wells. Then Classifier 1 is applied to the immune cell-only wells, where cell tracks are classified into two groups. The first group comprises cells that exhibit quick, irregular movement and change shape over time, identified as macrophages. The second group consists of motionless immune cells, and their cell type remains undetermined, for which we name them ‘Others’. **(2)** Myeloma and ‘Others’ single-cell tracks are similar and difficult to distinguish, for which we designed Classifier 2, which is specifically used to extract the features that can differentiate the two similar cell classes. Cell tracks from MM wells and the ‘Others’ single-cell tracks from immune cell-only wells form the dataset that is used to train Classifier 2.

After training, the LSTM-based classifiers can classify cells into three categories as is shown in Fig.4(b). The mix of three types of cells in a well makes classification a prerequisite for further analysis. Through classification, we can observe which cell type is responding to therapy, quantify each type, and determine the targeted effect of each immunotherapy on tumor/target cells.

### Benchmarking

In order to evaluate the performance of the detection and tracking methods, two contemporary methods are applied to the time-lapse cell images.

#### KTH-SE

KTH-SE [41] is a tracking-by-detection framework, which has shown strong performance across various datasets in Cell Tracking Challenges [56]. The detection method is based on threshold setting, and the tracking method utilizes the Viterbi algorithm. As this framework does not involve a machine learning method, there is no need for a training step. Furthermore, they offer a user-friendly graphical interface for ease of use.

#### YOLO

YOLO [44] is a CNN-based method comprising 24 convolutional layers. It utilizes a feature pyramid network to extract features from images. The output is a 7×7×30 vector representing 49 predictions, each comprising 2 bounding boxes and 20 class probabilities. YOLO has introduced its eighth version, which is faster, more accurate, and can detect and track more objects. In this study, YOLO v8 is employed for comparison with TLCellClassifier.

### Evaluation Metrics

To evaluate the performance of cell detection and tracking, this study utilizes metrics used for evaluating submissions in the Cell Tracking Challenge [57]. Both the detection and tracking methods undergo evaluation using the Acyclic Oriented Graph Matching (AOGM) measure. For evaluation purposes, two graphs are constructed: one for ground truth and one for prediction. In each graph, a node represents a detected cell, and edges denote one-to-one matches between two time points. This method assesses the difficulty of transforming the prediction graph into the ground truth graph, quantified as a weighted sum of the minimum graph operations required, including splitting, deleting, and adding a node, as well as deleting, adding, and altering the semantics of an edge, to align the two graphs. Numerically, DET is defined as a normalized AOGM measure for detection, with its value always falling within the [0,1] interval. A higher DET value indicates better performance. Similarly, TRA is defined as a normalized AOGM measure for tracking, also within the [0,1] range, where a higher value denotes better performance.

## Results

In this section, we evaluate the performance of the proposed framework, TLCellClassifier, by comparing it with KTH-SE and YOLO v8. The dataset consists of bright-field time-lapse images, each containing thousands of cells. Subsequent sections will assess the performance in detection, tracking, and classification.

### Cell Detection and Tracking

In this section, 640*×*640 images are cropped from the original 1328*×*1048 ones. Full-range images are not utilized here because the KTH-SE and YOLO methods cannot efficiently process such a large number of objects. The computational expense of the Viterbi algorithm makes it impractical for tracking thousands of objects, thus preventing the KTH-SE method from producing results for the 1328*×*1048 images within a few days on our server, equipped with an Intel Core i7-8700 processor and 32GB memory. The maximum resolution KTH-SE can handle for our dataset, which contains considerably fewer objects, is 640*×*640. Additionally, YOLO’s capability to detect objects is limited. In earlier versions of YOLO, it could detect a maximum of 49 objects. While subsequent versions of YOLO have significantly improved this number, our dataset presents a greater challenge due to the presence of thousands of cells in a single image, each with a low resolution of 12*×*12.

The comparison between TLCellClassifier and contemporary methods, KTH-SE and YOLO v8, is illustrated in Fig.5. In the bright-field images, there are over 600 cells. The ground truth for detection and tracking is manually annotated on these images. Both TLCellClassifier and KTH-SE demonstrate proficient detection capabilities, capturing most of the cells as illustrated in Fig.5(b) and (c). However, KTH-SE struggles to effectively differentiate between stromal cells and MM cells, as evident in Fig.5(c). Conversely, YOLO v8, depicted in Fig.5(d), detects only 300 cells, approximately half of the total count. This indicates YOLO’s limitation in detecting small cells, particularly in densely populated regions.

**Fig. 5.**
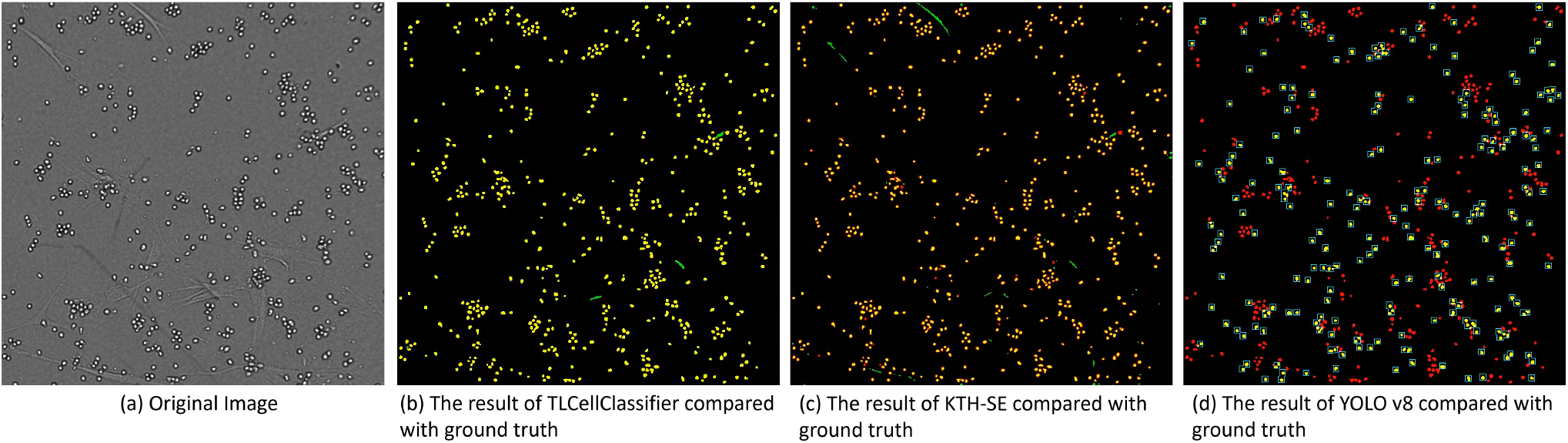
The comparison of the three methods: The ground truth segmentation is highlighted in red, while the predictions from the three methods are displayed in green. The overlap between the predicted region and the ground truth is represented in yellow (correct prediction) in (b) and (c). The boxes in (d) indicate the detection results of YOLO v8.

To conduct a comprehensive evaluation of cell detection and tracking accuracy of TLCellClassifier, ground truth from three wells is manually labeled. Subsequently, the accuracy of detection and tracking is assessed by comparing the predictions of the three methods against this ground truth. The evaluation metrics utilized in the Cell Tracking Challenge are employed, as described in the Evaluation Metrics subsection. The results of this assessment are calculated and presented in Fig.6. Notably, among the three methods, TLCellClassifier outperforms in both detection and tracking. Again, KTH-SE struggles to effectively differentiate stromal cells and MM cells, resulting in lower detection and tracking accuracy compared to TLCellClassifier. Consistent with the findings illustrated in Fig.5(d), YOLO v8 exhibits poor performance on small and crowded objects, thereby yielding the lowest detection and tracking accuracy among the three methods.

**Fig. 6.**
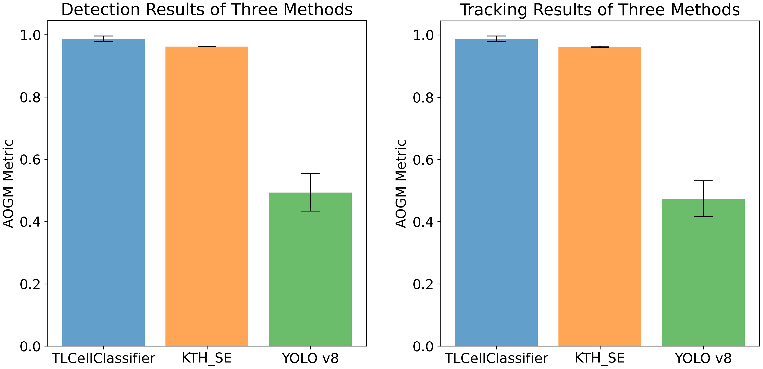
The comparison of TLCellClassifier, KTH-SE, and YOLO v8 on three wells: (a) cell detection performance; (b) cell tracking performance.

Based on the findings above, it’s evident that our dataset poses challenges beyond the capabilities of YOLO v8. To simplify the experiment, we cropped a smaller section of 480*×*480 from the full image. As illustrated in Fig.S1 in the Supplementary document, YOLO v8 successfully detects the majority of cells when the frame size is 480*×*480. However, with a frame size of 640*×*640, only half of the cells are detected.

The preceding section has outlined the detection and tracking results obtained for our dataset using the three methods. It is important to note that the images used above have a resolution of 640*×*640. Among these methods, only TLCellClassifier exhibits smooth performance even when the resolution is increased to 1328*×*1048.

### Cell Classification

It is important to categorize cells into different types: tumor (target), immune effector cells, and cells from the microenvironment, for accurate quantification of the treatment’s effect on a given patient’s tumor cells. To assess the performance of our LSTM classifier in the TLCellClassifier framework, we have prepared three different types of wells. The first type comprises cancer cell-only wells: 10 wells containing only MM cells. The second type consists of immune cell-only wells: 10 wells containing only CD14+ selected cells (macrophages/monocytes). In these wells, macrophages move rapidly and irregularly, while other cells (monocytes) remain relatively stable. The third type comprises mixed wells: 10 wells containing a mixture of cells from both types, where the number of immune cells is twice the number of cancer cells. The approach is implemented in several steps. Initially, cells are detected and tracked in time-lapse images. Subsequently, single-cell images (ROIs) are cropped to generate a track for each cell. The two-stage LSTM-based classification method is tested using two *in silico* experiments. In the first experiment, synthetic datasets with different ratios of each cell type are combined from the immune cell-only wells and cancer cell-only wells to evaluate the performance of the LSTM classifier. In the second experiment, the LSTM classifier is applied to the mixed-cell wells (i.e., the third type) to demonstrate the classification outcomes.

#### 1. Prediction of Different Cell Ratios

As mentioned earlier, single-cell tracks are extracted from immune cell-only wells or cancer cell-only wells based on cell detection and tracking. An LSTM-based binary classifier is then trained on the tracks from these two types of wells to discern the differences between the cells. Subsequently, the trained classifier is applied to the immune cell-only wells, where cell tracks are classified into two groups. The first group comprises cells that exhibit quick, irregular movement and change shape over time, identified as macrophages. The second group consists of cells that are motionless, and their cell type remains undetermined. Given three types of cells *−* myeloma cells, macrophages, and other cells (monocytes) *−* several synthetic datasets are created to classify cell tracks. The two-stage LSTM classifier is employed to classify these synthetic datasets. The results are illustrated in Fig.7.

**Fig. 7.**
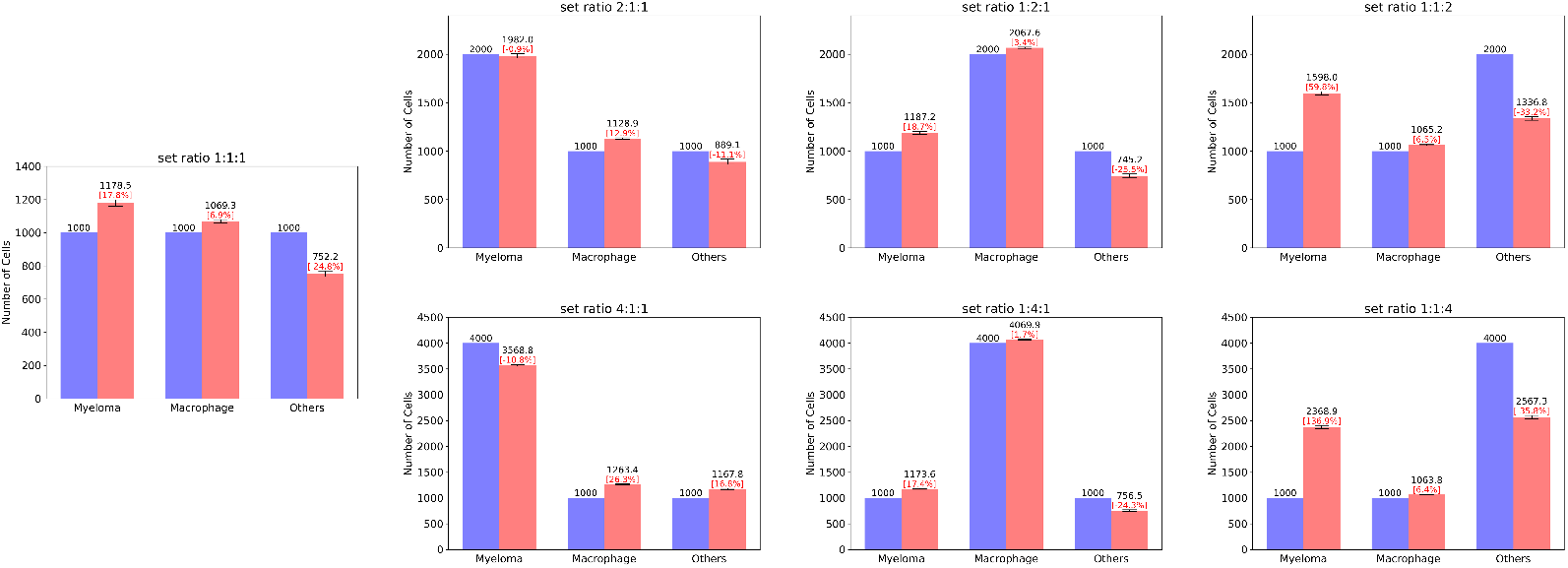
Classification of synthetic datasets. The experiment involves seven datasets, each containing different numbers of the three types of cells. The LSTM classifier is utilized to classify these cells into three categories. Each dataset is repeated 10 times, and the mean and standard deviation (in red text) of the predictions are calculated for each dataset. The blue bars represent the number of each cell type prepared in the dataset (ground truth), while the red bars depict the classification results.

In the figure, seven plots display the results of different *in silico* experiments using synthetic datasets as inputs. In each experiment, different numbers of the three cell types are prepared as ground truth, as indicated by the blue bars. Each of the seven experiments is repeated 10 times. The mean and standard deviation of the classification results are calculated and shown in the red bars. In the first column, an equal number of cells for each type (i.e., 1000 cells for each) is used, while the other three columns have a dominant cell type. It is observed that in the experiments represented in the left three columns, the accuracy is high. In contrast, the datasets in the rightmost column, which contain a higher proportion of ‘other’ cells (such as monocytes), exhibit relatively low classification accuracy. ‘Other’ cells, unlike macrophages which are actively moving, are relatively stable like myeloma cells. Thus, they are easily misclassified as myeloma cells when there are too many of them. Interestingly, the converse is not true; when a high number of myeloma cells are included, they are not misclassified as other cells at this image resolution level.

#### 2. Prediction on Mixed Cell Wells

In the preceding subsection, we evaluated the performance of the LSTM classifier using synthetic datasets with varying ratios of cell types. In this subsection, we rely on data from an actual *ex vivo* experiment, where twice the number of immune cells (macrophages/monocytes) are added in the well compared to myeloma cells at the beginning of the experiment. We have 10 wells with this condition, and we tested the two-stage LSTM classifier to estimate the proportion or number of myeloma cells relative to the immune cells. The results of this classification task are illustrated in Fig.8.

**Fig. 8.**
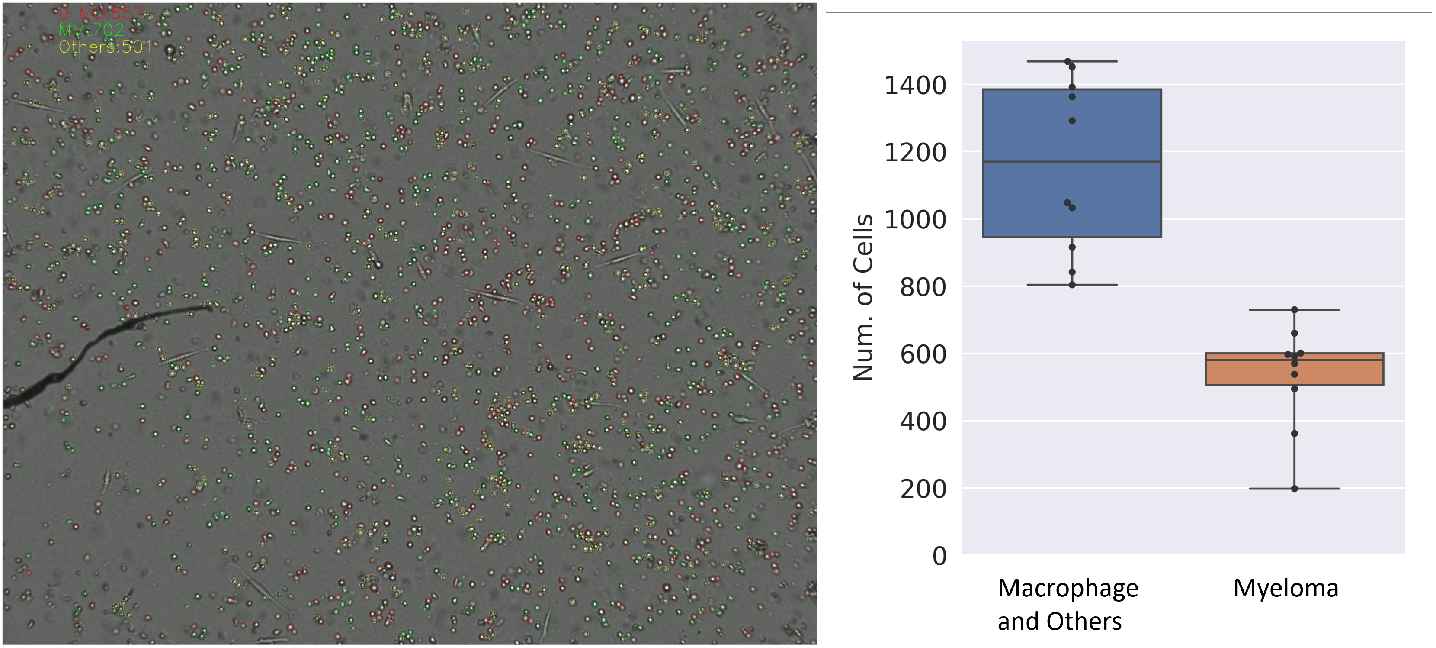
The left image displays a well containing three types of cells: macrophage (MO), myeloma (MM), and others. The cell classification is represented in three colors: red circles indicate macrophages (MO), green circles indicate myeloma (MM), and yellow circles indicate others. It is noted that the number of macrophages and others should be twice that of myeloma cells. The right plot illustrates the number of cells in each category at time point 0 across ten wells.

The left image in Fig.8 displays a well containing three types of cells: macrophage, myeloma, and others. Each cell type is visually distinguished with a specific color for classification. Macrophage cells are denoted by red circles, myeloma cells by green circles, and other cells by yellow circles. The right plot shows the classification results in ten wells. The number of cells in categories myeloma and immune cells (i.e., macrophage and others) is shown in the box plot. Given that the number of macrophages and others is twice the number of myeloma cells at the start, the right figure in Fig.8 shows the prediction is close to ground truth.

## Conclusion

Live cell imaging has been the subject of research in computer science and biomedical sciences for many years. Cells within the microenvironment pose a greater challenge for detection and tracking compared to general objects encountered in daily life. This paper presents TLCellClassifier, a framework designed to effectively detect, track, and classify cells. Operating on high-throughput time-lapse images characterized by low resolution and frame rates, TLCellClassifier achieves state-of-the-art performance. When compared to KTH-SE, a leading method in the Cell Tracking Challenge, and YOLO, renowned for general object detection, only TLCellClassifier demonstrates proficiency in both detection and tracking. Furthermore, it extends its applicability beyond cell tracking to encompass general object tracking, particularly for images featuring small objects such as aerial or fish images.

The cell type classification approach described in this article helps quantify the patient/immunotherapy-specific effect to profile patient tumors and immune effector cells to assess their efficacy prior to treatment. This ensures that every patient receives a therapeutic option to which they are most likely to respond, thus avoiding instances where patients have an underwhelming response. We submit that high-throughput *ex vivo* assays characterizing response to immunotherapies, when integrated with such specialized software, can help lay the groundwork for personalized medicine in cancer treatment.

## Funding

This work was supported by the National Science Foundation (NSF) [NSF-III2246796 and NSF-III2152030], and the Pentecost Family Foundation and Pentecost Family Myeloma Research Center (PMRC) at Moffitt Cancer Center. It was also supported in part by the H. Lee Moffitt Cancer Center Physical Sciences in Oncology (PSOC) Grant 1U54CA193489-01A1 (to K.H.S., A.S.S., R.J.G. and R.A.G.), H. Lee Moffitt Cancer Center’s Team Science Grant (to A.S.S. and K.H.S.), Miles for Moffitt Foundation (to A.S.S.) and the Cancer Center Support Grant P30-CA076292 to the Moffitt Cancer Center. A

## Notes

### Competing Interest Statement

The authors have declared no competing interest.

## References

1. Ira Mellman, George Coukos, and Glenn Dranoff. Cancer immunotherapy comes of age. Nature, 480(7378):480–489, 2011.

2. Yiping Yang et al. Cancer immunotherapy: harnessing the immune system to battle cancer. The Journal of clinical investigation, 125(9):3335–3337, 2015.

3. Stephen J Till, James N Francis, Kayhan Nouri-Aria, and Stephen R Durham. Mechanisms of immunotherapy. Journal of Allergy and Clinical Immunology, 113(6):1025– 1034, 2004.

4. Manfred Schuster, Andreas Nechansky, and Ralf Kircheis. Cancer immunotherapy. Biotechnology Journal: Healthcare Nutrition Technology, 1(2):138–147, 2006.

5. James C Yang, Marybeth Hughes, Udai Kammula, Richard Royal, Richard M Sherry, Suzanne L Topalian, Kimberly B Suri, Catherine Levy, Tamika Allen, Sharon Mavroukakis, et al. Ipilimumab (anti-ctla4 antibody) causes regression of metastatic renal cell cancer associated with enteritis and hypophysitis. Journal of immunotherapy, 30(8):825–830, 2007.

6. Apostolia-Maria Tsimberidou. Targeted therapy in cancer. Cancer chemotherapy and pharmacology, 76:1113–1132, 2015.

7. Zayar P Khin, Maria LC Ribeiro, Timothy Jacobson, Lori Hazlehurst, Lia Perez, Rachid Baz, Kenneth Shain, and Ariosto S Silva. A preclinical assay for chemosensitivity in multiple myeloma. Cancer research, 74(1):56–67, 2014.

8. Ariosto Silva, Timothy Jacobson, Mark Meads, Allison Distler, and Kenneth Shain. An organotypic high throughput system for characterization of drug sensitivity of primary multiple myeloma cells. JoVE (Journal of Visualized Experiments), (101):e53070, 2015.

9. Ariosto Silva, Maria C Silva, Praneeth Sudalagunta, Allison Distler, Timothy Jacobson, Aunshka Collins, Tuan Nguyen, Jinming Song, Dung-Tsa Chen, Lu Chen, et al. An ex vivo platform for the prediction of clinical response in multiple myeloma. Cancer research, 77(12):3336–3351, 2017.

10. Praneeth Sudalagunta, Maria C Silva, Rafael R Canevarolo, Raghunandan Reddy Alugubelli, Gabriel DeAvila, Alexandre Tungesvik, Lia Perez, Robert Gatenby, Robert Gillies, Rachid Baz, et al. A pharmacodynamic model of clinical synergy in multiple myeloma. EBioMedicine, 54:102716, 2020.

11. Hao Wu, Jovial Niyogisubizo, Keliang Zhao, Jintao Meng, Wenhui Xi, Hongchang Li, Yi Pan, and Yanjie Wei. A weakly supervised learning method for cell detection and tracking using incomplete initial annotations. International Journal of Molecular Sciences, 24(22):16028, 2023.

12. Qibing Jiang, Praneeth Sudalagunta, Maria C Silva, Rafael R Canevarolo, Xiaohong Zhao, Khandakar Tanvir Ahmed, Raghunandan Reddy Alugubelli, Gabriel DeAvila, Alexandre Tungesvik, Lia Perez, et al. Cancercelltracker: a brightfield time-lapse microscopy framework for cancer drug sensitivity estimation. Bioinformatics, 38(16):4002–4010, 2022.

13. W Joost Lesterhuis, John BAG Haanen, and Cornelis JA Punt. Cancer immunotherapy–revisited. Nature reviews Drug discovery, 10(8):591–600, 2011.

14. Frits Zernike. Phase contrast, a new method for the microscopic observation of transparent objects part ii. Physica, 9(10):974–986, 1942.

15. RD Allen and GB David. The zeiss-nomarski differential interference equipment for transmitted-light microscopy. Zeitschrift fur wissenschaftliche Mikroskopie und mikroskopische Technik, 69(4):193–221, 1969.

16. Xavier Michalet, Achillefs N Kapanidis, Ted Laurence, Fabien Pinaud, Soeren Doose, Malte Pflughoefft, and Shimon Weiss. The power and prospects of fluorescence microscopies and spectroscopies. Annual review of biophysics and biomolecular structure, 32(1):161–182, 2003.

17. Vladimír Ulman, Martin Maška, Klas EG Magnusson, Olaf Ronneberger, Carsten Haubold, Nathalie Harder, Pavel Matula, Petr Matula, David Svoboda, Miroslav Radojevic, et al. An objective comparison of cell-tracking algorithms. Nature methods, 14(12):1141–1152, 2017.

18. Christopher J Soelistyo, Kristina Ulicna, and Alan R Lowe. Machine learning enhanced cell tracking. Frontiers in Bioinformatics, 3, 2023.

19. Junya Hayashida and Ryoma Bise. Cell tracking with deep learning for cell detection and motion estimation in low-frame-rate. In Medical Image Computing and Computer Assisted Intervention–MICCAI 2019: 22nd International Conference, Shenzhen, China, October 13–17, 2019, Proceedings, Part I 22, pages 397–405. Springer, 2019.

20. Tomas Vicar, Jan Balvan, Josef Jaros, Florian Jug, Radim Kolar, Michal Masarik, and Jaromir Gumulec. Cell segmentation methods for label-free contrast microscopy: review and comprehensive comparison. BMC bioinformatics, 20:1–25, 2019.

21. Felix Buggenthin, Carsten Marr, Michael Schwarzfischer, Philipp S Hoppe, Oliver Hilsenbeck, Timm Schroeder, and Fabian J Theis. An automatic method for robust and fast cell detection in bright field images from high-throughput microscopy. BMC bioinformatics, 14(1):297, 2013.

22. Shutong Tse, Laura Bradbury, Justin WL Wan, Haig Djambazian, Robert Sladek, and Thomas Hudson. A combined watershed and level set method for segmentation of brightfield cell images. In Medical Imaging 2009: Image Processing, volume 7259, pages 1147–1156. SPIE, 2009.

23. Fabian Isensee, Paul F Jäger, Simon AA Kohl, Jens Petersen, and Klaus H Maier-Hein. Automated design of deep learning methods for biomedical image segmentation. arXiv preprint arXiv:1904.08128, 2019.

24. Olaf Ronneberger, Philipp Fischer, and Thomas Brox. U-net: Convolutional networks for biomedical image segmentation. In International Conference on Medical image computing and computer-assisted intervention, pages 234–241. Springer, 2015.

25. Yanming Zhu and Erik Meijering. Automatic improvement of deep learning-based cell segmentation in time-lapse microscopy by neural architecture search. Bioinformatics, 37(24):4844–4850, 2021.

26. Alexandre Dufour, Roman Thibeaux, Elisabeth Labruyere, Nancy Guillen, and Jean-Christophe Olivo-Marin. 3-d active meshes: fast discrete deformable models for cell tracking in 3-d time-lapse microscopy. IEEE transactions on image processing, 20(7):1925–1937, 2010.

27. Martin Maška, Ondřej Daněk, Saray Garasa, Ana Rouzaut, Arrate Munoz-Barrutia, and Carlos Ortiz-de Solorzano. Segmentation and shape tracking of whole fluorescent cells based on the chan–vese model. IEEE transactions on medical imaging, 32(6):995–1006, 2013.

28. Oleh Dzyubachyk, Wiggert A Van Cappellen, Jeroen Essers, Wiro J Niessen, and Erik Meijering. Advanced level-set-based cell tracking in time-lapse fluorescence microscopy. IEEE transactions on medical imaging, 29(3):852–867, 2010.

29. Alexandre Dufour, Vasily Shinin, Shahragim Tajbakhsh, Nancy Guillén-Aghion, J-C Olivo-Marin, and Christophe Zimmer. Segmenting and tracking fluorescent cells in dynamic 3-d microscopy with coupled active surfaces. IEEE Transactions on Image Processing, 14(9):1396–1410, 2005.

30. Harold W Kuhn. The hungarian method for the assignment problem. Naval research logistics quarterly, 2(1-2):83–97, 1955.

31. Erick Moen, Enrico Borba, Geneva Miller, Morgan Schwartz, Dylan Bannon, Nora Koe, Isabella Camplisson, Daniel Kyme, Cole Pavelchek, Tyler Price, et al. Accurate cell tracking and lineage construction in live-cell imaging experiments with deep learning. Biorxiv, page 803205, 2019.

32. Qibing Jiang, Praneeth Sudalagunta, Mark B Meads, Khandakar Tanvir Ahmed, Tara Rutkowski, Ken Shain, Ariosto S Silva, and Wei Zhang. An Advanced Framework for Time-lapse Microscopy Image Analysis. bioRxiv, 2020.

33. Klas EG Magnusson, Joakim Jaldén, Penney M Gilbert, and Helen M Blau. Global linking of cell tracks using the viterbi algorithm. IEEE transactions on medical imaging, 34(4):911–929, 2014.

34. Engin Türetken, Xinchao Wang, Carlos J Becker, Carsten Haubold, and Pascal Fua. Network flow integer programming to track elliptical cells in time-lapse sequences. IEEE transactions on medical imaging, 36(4):942–951, 2016.

35. Ryoma Bise, Zhaozheng Yin, and Takeo Kanade. Reliable cell tracking by global data association. In 2011 IEEE International Symposium on Biomedical Imaging: From Nano to Macro, pages 1004–1010. IEEE, 2011.

36. Na Dong, Li Zhao, Chun-Ho Wu, and Jian-Fang Chang. Inception v3 based cervical cell classification combined with artificially extracted features. Applied Soft Computing, 93:106311, 2020.

37. César Cheuque, Marvin Querales, Roberto León, Rodrigo Salas, and Romina Torres. An efficient multi-level convolutional neural network approach for white blood cells classification. Diagnostics, 12(2):248, 2022.

38. Chao Liu, Zhuo Wang, Junkun Jia, Qiao Liu, and Xuantao Su. High-content video flow cytometry with digital cell filtering for label-free cell classification by machine learning. Cytometry Part A, 103(4):325–334, 2023.

39. Hongda Wang, Hatice Ceylan Koydemir, Yunzhe Qiu, Bijie Bai, Yibo Zhang, Yiyin Jin, Sabiha Tok, Enis Cagatay Yilmaz, Esin Gumustekin, Yair Rivenson, et al. Early detection and classification of live bacteria using time-lapse coherent imaging and deep learning. Light: Science & Applications, 9(1):118, 2020.

40. Colton Boudreau, Tse-Luen Wee, Yan-Rung Duh, Melissa P Couto, Kimya H Ardakani, and Claire M Brown. Excitation light dose engineering to reduce photo-bleaching and phototoxicity. Scientific reports, 6(1):30892, 2016.

41. Klas E. G. Magnusson, Joakim Jaldén, Penney M. Gilbert, and Helen M. Blau. Global linking of cell tracks using the viterbi algorithm. IEEE Transactions on Medical Imaging, 34(4):911–929, 2015.

42. Kunihiko Fukushima. Neocognitron: A self-organizing neural network model for a mechanism of pattern recognition unaffected by shift in position. Biological cybernetics, 36(4):193–202, 1980.

43. Kaiming He, Xiangyu Zhang, Shaoqing Ren, and Jian Sun. Deep residual learning for image recognition. In Proceedings of the IEEE conference on computer vision and pattern recognition, pages 770–778, 2016.

44. Joseph Redmon, Santosh Divvala, Ross Girshick, and Ali Farhadi. You only look once: Unified, real-time object detection. In Proceedings of the IEEE conference on computer vision and pattern recognition, pages 779–788, 2016.

45. Christian Szegedy, Wei Liu, Yangqing Jia, Pierre Sermanet, Scott Reed, Dragomir Anguelov, Dumitru Erhan, Vincent Vanhoucke, and Andrew Rabinovich. Going deeper with convolutions. In Proceedings of the IEEE conference on computer vision and pattern recognition, pages 1–9, 2015.

46. Mengzi Hu, Ziyang Li, Jiong Yu, Xueqiang Wan, Haotian Tan, and Zeyu Lin. Efficient-lightweight yolo: improving small object detection in yolo for aerial images. Sensors, 23(14):6423, 2023.

47. Yang Liu, Peng Sun, Nickolas Wergeles, and Yi Shang. A survey and performance evaluation of deep learning methods for small object detection. Expert Systems with Applications, 172:114602, 2021.

48. Kang Tong, Yiquan Wu, and Fei Zhou. Recent advances in small object detection based on deep learning: A review. Image and Vision Computing, 97:103910, 2020.

49. Fatih Cagatay Akyon, Sinan Onur Altinuc, and Alptekin Temizel. Slicing aided hyper inference and fine-tuning for small object detection. In 2022 IEEE International Conference on Image Processing (ICIP), pages 966–970. IEEE, 2022.

50. Liu Yang and Rong Jin. Distance metric learning: A comprehensive survey. Michigan State Universiy, 2(2):4, 2006.

51. Long Short-Term Memory. Long short-term memory. Neural computation, 9(8):1735–1780, 2010.

52. Nicolai Wojke and Alex Bewley. Deep cosine metric learning for person re-identification. In 2018 IEEE winter conference on applications of computer vision (WACV), pages 748–756. IEEE, 2018.

53. Nicolai Wojke, Alex Bewley, and Dietrich Paulus. Simple online and realtime tracking with a deep association metric. In 2017 IEEE international conference on image processing (ICIP), pages 3645–3649. IEEE, 2017.

54. Christian Szegedy, Vincent Vanhoucke, Sergey Ioffe, Jon Shlens, and Zbigniew Wojna. Rethinking the inception architecture for computer vision. In Proceedings of the IEEE conference on computer vision and pattern recognition, pages 2818–2826, 2016.

55. Jia Deng, Wei Dong, Richard Socher, Li-Jia Li, Kai Li, and Li Fei-Fei. Imagenet: A large-scale hierarchical image database. In 2009 IEEE conference on computer vision and pattern recognition, pages 248–255. Ieee, 2009.

56. Martin Maška, Vladimír Ulman, Pablo Delgado-Rodriguez, Estibaliz Gómez-de Mariscal, Tereza Nečasová, Fidel A Guerrero Peña, Tsang Ing Ren, Elliot M Meyerowitz, Tim Scherr, Katharina Löffler, et al. The cell tracking challenge: 10 years of objective benchmarking. Nature Methods, pages 1–11, 2023.

57. Pavel Matula, Martin Maška, Dmitry V Sorokin, Petr Matula, Carlos Ortiz-de Solórzano, and Michal Kozubek. Cell tracking accuracy measurement based on comparison of acyclic oriented graphs. PloS one, 10(12):e0144959, 2015.

