## Supplementary material for "TLCellClassifier: Machine Learning Based Cell Classification for Bright-Field Time-Lapse Images": Fig.S1

First Author

### 1 The evaluation of YOLO v8 on different size of images

Our research work demonstrates that our dataset presents challenges beyond the capabilities of YOLO v8 [1]. To simplify the experiment, we cropped a smaller section of 480×480 from the full image. As illustrated in Fig.S1, YOLO v8 successfully detects the majority of cells when the frame size is 480×480. However, with a frame size of 640×640, only half of the cells are detected.

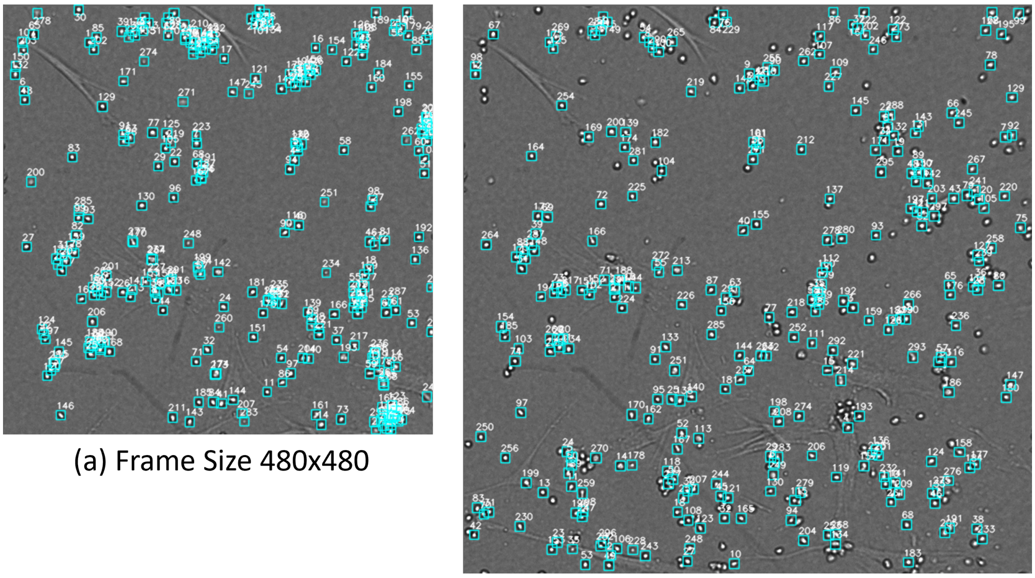

(a) Frame Size 480x480

(b) Frame Size 640x640

Fig. S1 Explore the performance of YOLO v8: (a) YOLO v8 can detect 94.62% of cells when the frame size is 480×480. (b) YOLO v8 can only detect 54.26% of cells when the frame size is 640×640.

### References

[1] Joseph Redmon, Santosh Divvala, Ross Girshick, and Ali Farhadi. You only look once: Unified, real-time object detection. In *Proceedings of the IEEE conference on computer vision and pattern recognition*, pages 779–788, 2016.
